# Impact of Image Gently Campaign at a Medium-Sized Teaching Hospital

**DOI:** 10.1101/057265

**Authors:** Eric Liu, Yen-Ying Wu

## Introduction

More than 62 million computed tomography (CT) scans are performed each year in the United States, compared with 3 million in 1980. Approximately 4 million CT scans per year are performed on children.^1^ Recent literature suggests that approximately 2% of future cancers may be directly attributable to CT scanning in children. One study estimated 4,350 future cancers could be induced by 1 year of pediatric CT imaging in the United States.^2^ Pediatric patients are at higher risk for developing cancer because they are more radiosensitive and have longer postradiation life spans than adults. The Society for Pediatric Radiology implemented the Image Gently Campaign on January 22, 2008 to raise awareness and promote methods to reduce radiation dose.^3^ Methods to reduce radiation dose in children include utilizing optimized ionizing imaging when necessary and utilizing nonionizing imaging alternatives when possible.^4^

We sought to determine if the imaging of pediatric patients at Loma Linda University Medical Center (LLUMC) was affected since the Image Gently Campaign. Specifically, we reviewed the trend of the number of ionizing exams before and after the implementation of the Image Gently Campaign, and if there has been a shift from the use of CT as the initial imaging examination in pediatric patients presenting to the emergency department with suspected appendicitis.

## Materials and Methods

An institutional review board approved retrospective study was performed on the LLUMC picture archiving and communication system (PACS). The number of CT, radiograph, ultrasound (US), magnetic resonance imaging (MRI), and total number of imaging exams performed in pediatric patients (18 years of age and under), were determined for each year between 2003 and 2013.

For pediatric patients from the emergency department with suspected appendicitis, a retrospective search was performed on PACS and the imaging algorithm was investigated. This was done for years 2007, 2008, and 2013. For these patients, the imaging algorithm was investigated by determining whether the patient received an US or CT exam first.

## Results

Between 2003 and 2013, the number of non-ionizing imaging exams increased while the number of ionizing imaging exams decreased. The total imaging examination peak was in 2005 (73,010 exams). The mean total number of exams prior to the Image Gently Campaign (2003-2007) was 69,796. The mean total number of non-ionizing exams from 2003-2007 was 9,987. The mean total number of ionizing exams from 2003-2007 was 59,809. Comparing the data from 2013 with the mean from 2003-2007, there was a 12% decrease in the total number of imaging exams, a 16% increase in the number of non-ionizing imaging exams and a 21% decrease inthe number of ionizing imaging exams(**Figure 1**).

**Fig. 1.**
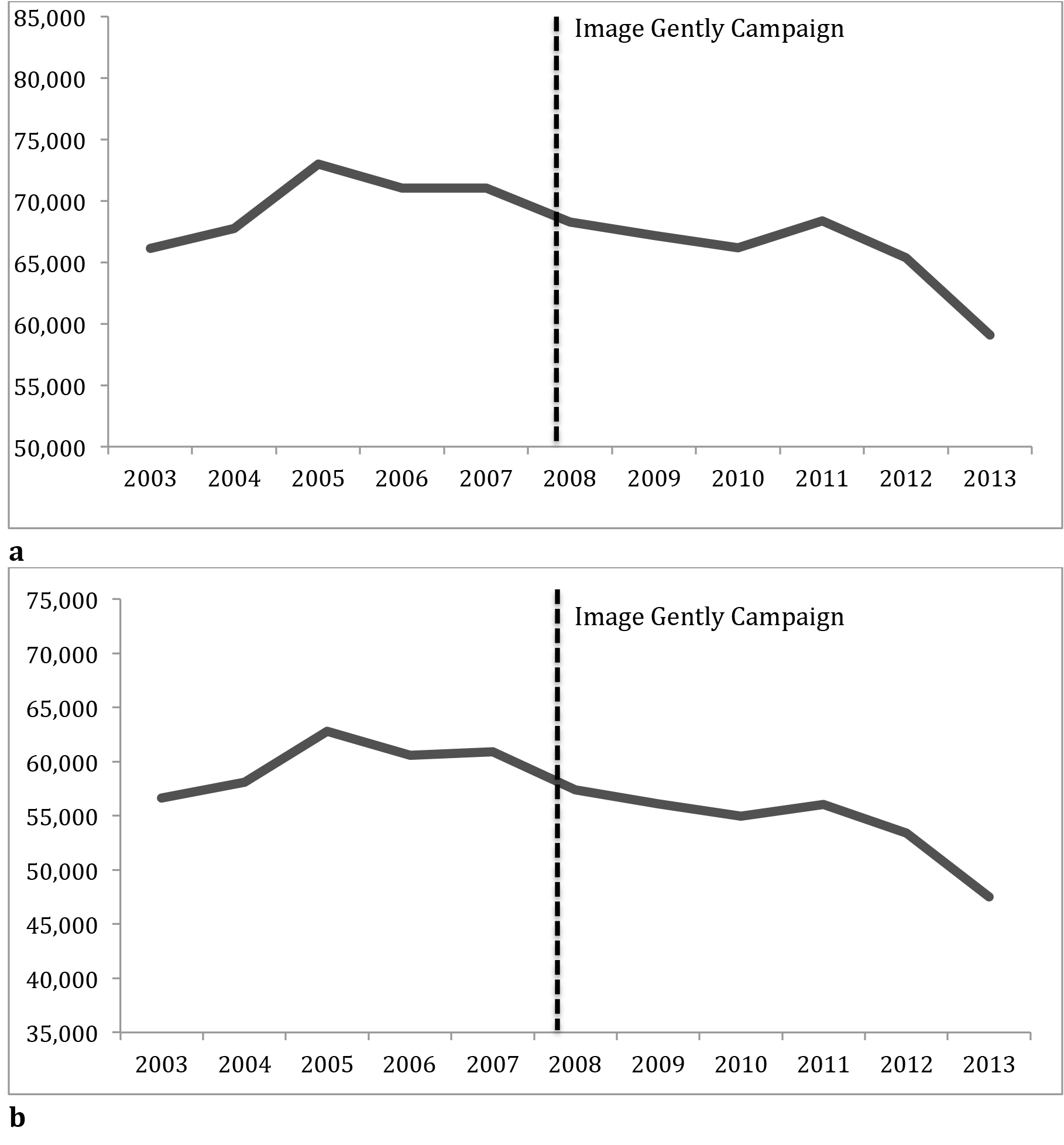
Line graph representation of the trend of the number of imaging exams before andafter the Image Gently Campaign in 2008. (**a**) Overall total imaging exams. (**b**) Total ionizing exams, (**c**) Total non-ionizing exams.

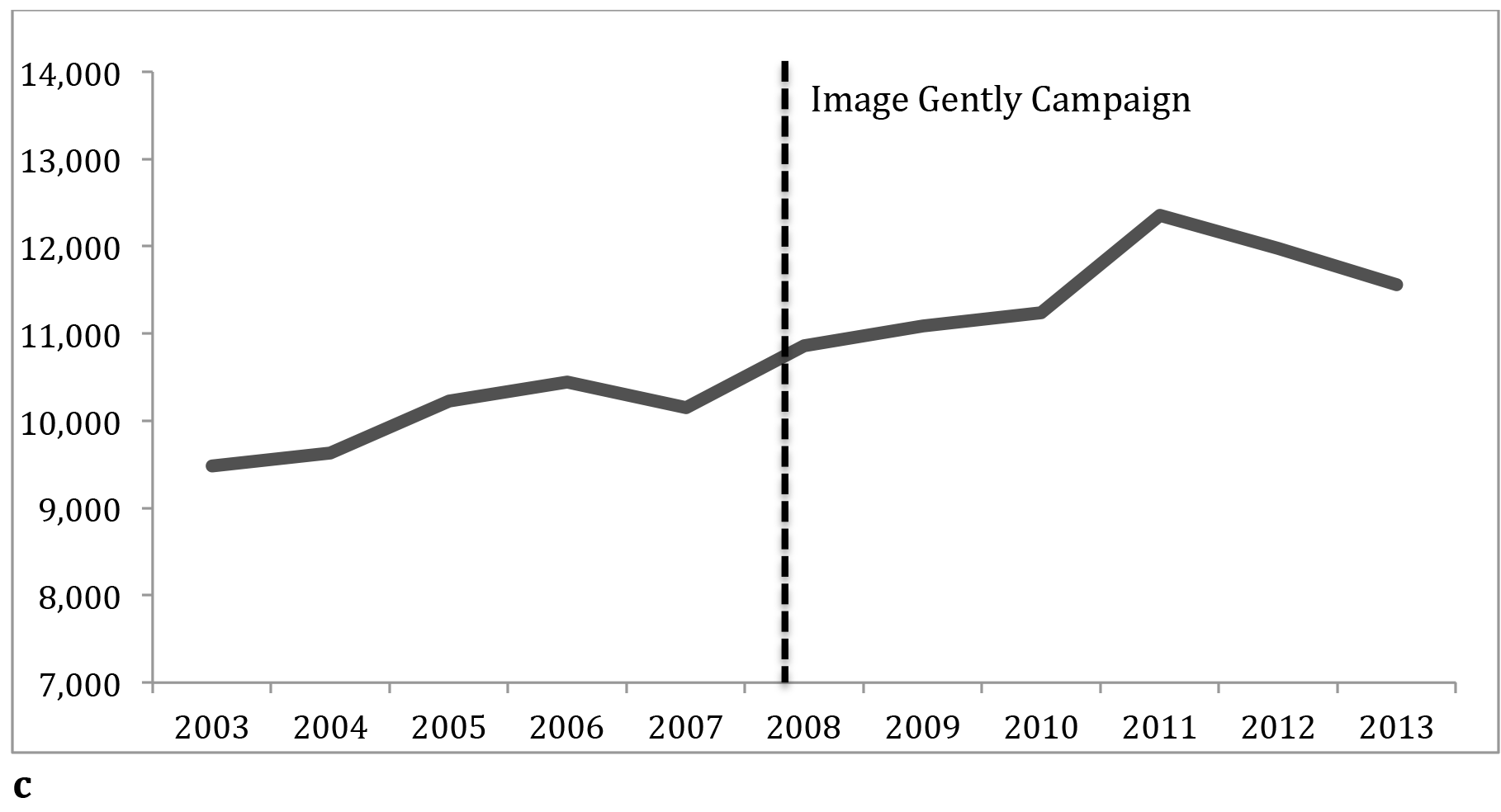

The total number of imaging exams have decreased largely due to a decrease in CT and radiograph exams while there has been an increase in the number of MRI and US exams (**Figure 2**). Radiograph exams account for the majority of ionizing imaging exams. The mean total number of radiograph exams prior to 2008 was 52,492. The mean total number of CT exams prior to 2008 was 7,317. The mean total number of MRI exams prior to the 2008 was 3,892. The mean total number of US exams prior to 2008 was 6,095. When comparing the mean prior to the Image Gently Campaign to the data from 2013, there has been a 33% decrease in the number of CT exams, a 19% decrease in the number of radiograph exams, a 13% increase in the number of US exams, and a 19% increase in the number of MRI exams. The mean total number of CT abdomen and pelvis exams prior to the Image Gently Campaign was 2,247. For CT abdomen and pelvis exams, there was a 64% decrease in 2013 from the mean prior to 2008 (**Figure 3**).

**Fig. 2.**
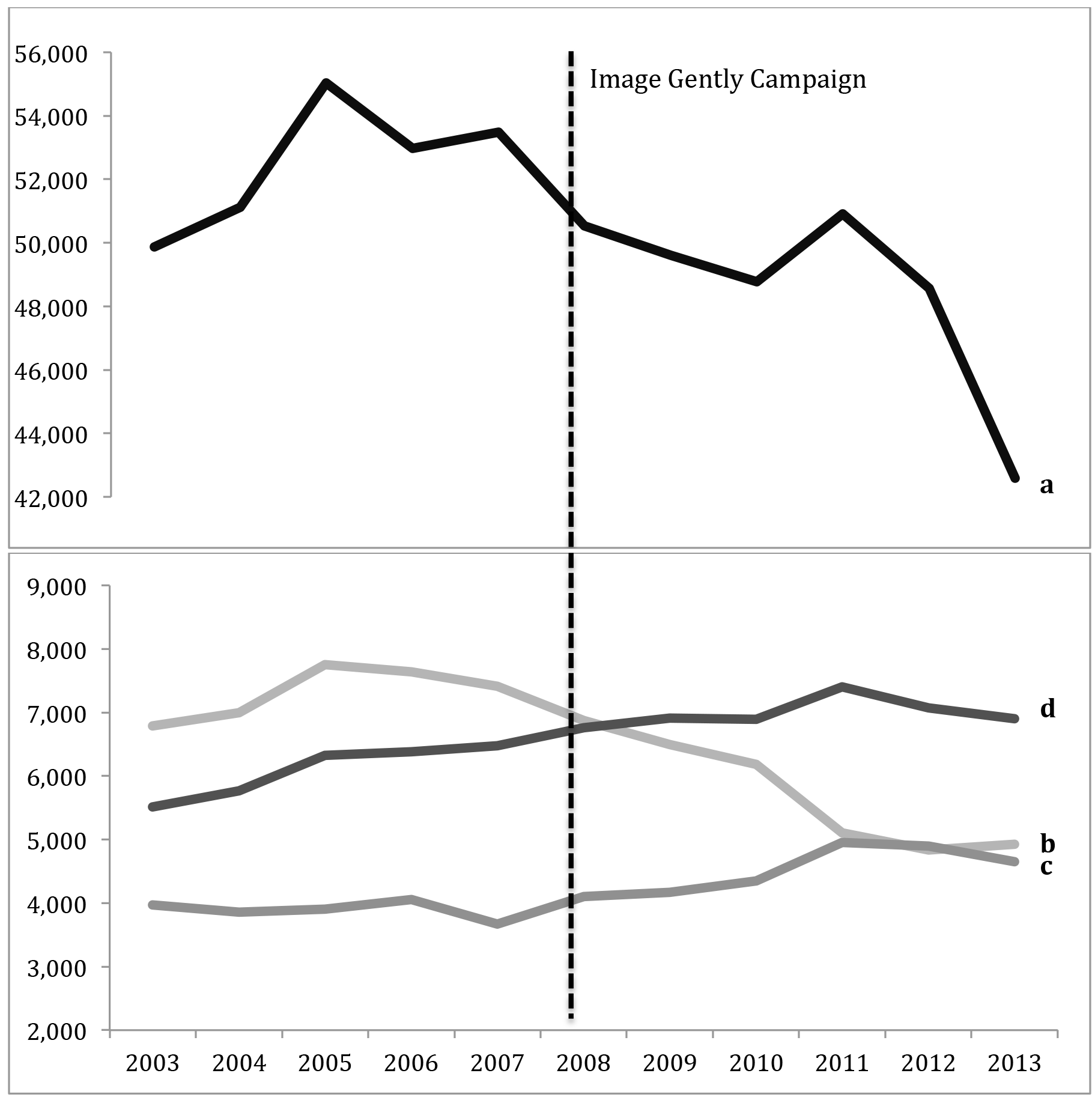
Line graph representation of the trend of the number of exams before and after the Image Gently Campaign in 2008. (a) Radiograph exams. (b) CT exams. (c) MRI exams. (d) US exams.

**Fig. 3.**
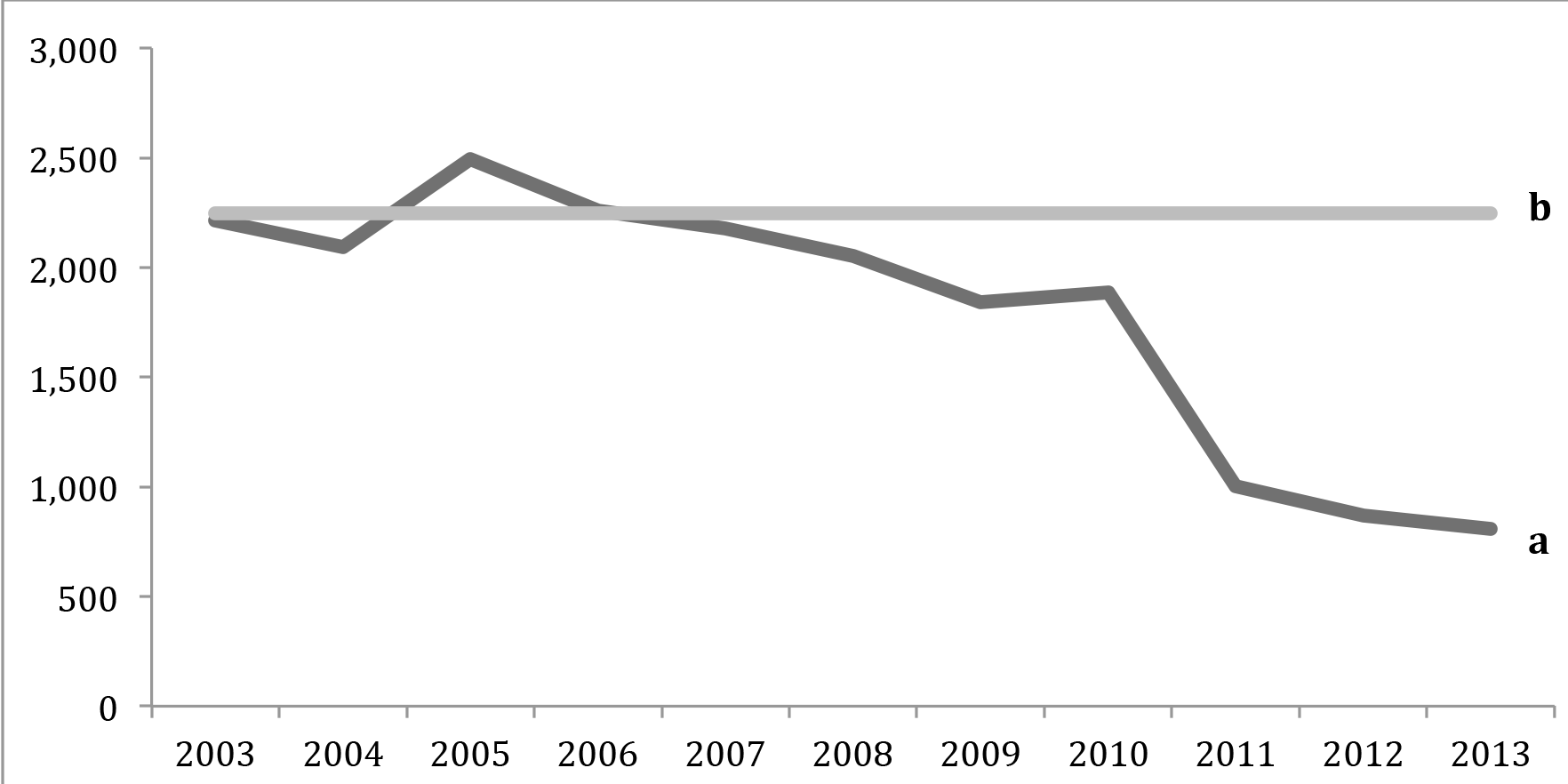
Line graph representation of the trend of the total number of CT abdomen and pelvis exams (**a**) with mean CT abdomen and pelvis exams prior to the Image Gently Campaign (**b**).

In 2007, 37.5% of patients received an US and 62.5% received a CT as the initial examination for suspected appendicitis. In 2008, the initial exam was US 50% and CT 50%. In 2013, the initial exam was US 85.4% and CT 14.6%.

## Discussion

### Overall reversed trend of ionizing and non-ionizing imaging

Since the implementation of the Image Gently Campaign in 2008, the number of ionizing imaging exams has decreased while non-ionizing imaging exam alternatives has increased at LLUMC. When comparing the mean prior to the Image Gently Campaign (2003-2007) to data from 2013, there has been a 33% decrease in the number of CT exams and a 19% decrease in the number of radiograph exams. Conversely, there has been a 13% increase in the number of US exams and a 19% increase in the number of MRI exams.

This is in-line with the previously published results from the 2013 retrospective study by Miglioretti et al. that reviewed the use of CT in children younger than 15 years of age from 1996 to 2010 across seven US health care systems, which included Kaiser Permanente and the Henry Ford Health System. The study found that the use of CT on older children nearly tripled from 1996 to 2005 to a peak of 27 CT scans per 1000 children. The use of CT in the study population stabilized and slightly declined since 2007, particularly among younger children.

The study attributed the decline to the increased awareness about the cancer risksfrom pediatric imaging in part due to the Image Gently Campaign. The decline in CT use in pediatric patients at LLUMC parallels the trend across the country.^5^

### CT abdomen and pelvis work-up shift

The risk of radiation-induced solid cancer is highest for CT scans of the abdomen and pelvis, which has seen the most dramatic increase in use, especially among older children.^5^ At LLUMC, the number of CT abdomen and pelvis exams has decreased by 64% in 2013 from the mean prior to the Image Gently Campaign. The Miglioretti et al. study found that most of the CT abdomen and pelvis exams were done for pain (40%), appendicitis (11%) or infection (6%). CT use in pediatric patients can be limited to those with equivocal or negative findings on US.^6^ Our focused review of the years 2007, 2008, and 2013 confirms a decreased utilization of CT and an increased utilization of US as the initial exam in the imaging work-up for appendicitis.

### Study Limitations

There are reasons that may explain the reduction of ionizing radiation exams other than the Image Gently Campaign. Possible patient population reduction or a downturn in the economy was not correlated. However, these reasons do not have the same obvious effects on nonionizing exams, which have increased. A viable explanation to account for the increased use of non-ionizing imaging alternatives is the technological improvements and availability of modalities such as US and MRI. Change in reimbursement is another potential reason for the observed increased utilization of non-ionizing exams since 2008.

The study explores exam frequency and not actual dose reduction. Given improvements in CTand radiograph technology and technique, actual dose is likely to have reduced as well. Non-ionizing imaging alternatives are not without their disadvantages. Cost, need for sedation, and differences in sensitivity/specificity were not reviewed in this study.

## Conclusions

Since the Imaging Gently Campaign, there has been a shift from the use of ionizing imaging exams in pediatric patients in favor of non-ionizing imaging alternatives such as US and MRI.

## References

1 Smith-Bindman R, Miglioretti DL, Johnson E, et al. Use of diagnostic imaging studies and associated radiation exposure for patients enrolled in large integrated health care systems, 1996-2010. JAMA. 2012; 307(22): 2400–2409.

2 Berrington de Gonzalez A, Mahesh M, Kim KP, et al. Projected cancer risks from computed tomographic scans performed in the United States in 2007. Arch Intern Med. 2009; 169(22): 2071–2077.

3 Image Gently: The Alliance for Radiation Safety in Pediatric Imaging. CT-what can I do? http://pedrad.org/associations/5364/ig/index.cfm?page=368 Accessed November 29, 2014.

4 Shah, NB, Platt SL. ALARA: Is there a cause for alarm? Reducing radiation risks from computed tomography scanning in children. Curr Opin Pediatr. 2008; 20(3): 243–247.

5 Miglioretti DL, Johnson E, Williams A, et al. The use of computed tomography in pediatrics and the associated radiation exposure and estimated cancer risk. JAMA Pediatrics. 2013: 167(8): 700–707.

6 Garcia Pena BM, Mandl KD, Kraus SJ, et al. Ultrasonography and limited computed tomography in the diagnosis and management of appendicitis in children. JAMA. 1999; 282(11): 1041–1046.

